# Molecular basis of BET family protein inhibition by four clinical-stage small-molecule inhibitor drugs

**DOI:** 10.64898/2026.02.05.704030

**Authors:** Sheng Lin, Yu You, Xiaofeng Chen, Guangwen Lu

## Abstract

The bromodomain and extra-terminal (BET) family proteins are critical epigenetic readers involved in various diseases, making them attractive therapeutic targets. This study elucidates the molecular basis of inhibition by four clinical-stage BET inhibitors (ZEN-3694, INCB054329, PLX51107, and INCB057643) through detailed structural comparison and analysis. Using X-ray crystallography, we determined their complex structures with the second bromodomain of human BRD2 at 1.4-1.5 Å resolution. All inhibitors occupy the conserved acetyl-lysine binding pocket via extensive hydrophobic interactions. This hydrophobic network is further coupled with hydrogen-bond contacts for inhibitor-binding stabilization. A common feature is the formation of at least one hydrogen bond with a conserved residue of N429. Notably, INCB057643 exhibits an enhanced polar network, forming additional hydrogen bonds with residues D377 and N429, which may contribute to its improved pharmacological profile. These structures reveal a shared competitive inhibition mechanism by occluding the acetyl-lysine binding site, providing a rational basis for their transcriptional repression and guiding future inhibitor design.

## Main text

Bromodomain-containing proteins (BRDs) are evolutionarily conserved protein modules that function as specific readers of ε-N-lysine acetylation marks (1), a fundamental post-translational modification implicated in chromatin remodeling and transcriptional regulation. The bromodomains achieve their biological function by recognizing and binding to acetylated lysine (Kac) residues on histone tails and various non-histone proteins. In humans, 46 distinct bromodomain-containing proteins have been identified and systematically classified into eight major families (2). Among these, the bromodomain and extra-terminal (BET) family represents a critically important subclass due to its profound involvement in the occurrence and progression of a wide spectrum of human diseases. The BET family’s role spans numerous cancers (including but not limited to pancreatic ductal adenocarcinoma, acute lymphoblastic leukemia, and breast cancer) (3), cardiovascular pathologies (such as hypertension and atherosclerotic plaque formation) (4), and the dysregulation of inflammatory gene transcription cascades (5). The human BET family comprises four members: BRD2, BRD3, BRD4, and the testis-specific protein BRDT. Each shares a conserved architecture featuring two N-terminal bromodomains (BD1 and BD2, each approximately 110 amino acids) and a unique C-terminal extra-terminal (ET) domain. This structural design empowers BET proteins to act as pivotal epigenetic scaffolds and regulators of gene expression, orchestrating essential cellular processes including proliferation, cell cycle progression, and programmed cell death.

Notably, aberrant expression and function of BRD2 have been directly linked to oncogenesis and other disorders (6, 7). For instance, its overexpression in leukemias and lymphomas drives the transcription of key oncogenes (6), whereas reduced BRD2 levels in animal models correlate with an increased susceptibility to seizures (7). Given the central role of BET proteins in disease, the development of small-molecule inhibitors that disrupt their function has emerged as a highly promising therapeutic strategy, particularly in oncology. Consequently, BET inhibitors constitute a major focal point in anticancer drug discovery. A significant number of candidate molecules have entered clinical evaluation, including ABBV-744, SJ432, RVX-208, ZEN-3694, INCB054329, INCB057643, PLX51107, and etc. Among these, four compounds— ZEN-3694 (8) (NCT03901469), INCB054329 (9) (NCT02431260), INCB057643 (10) (NCT02711137), and PLX51107 (11) (NCT02683395)—have advanced to Phase II clinical trials. These inhibitors are designed primarily to target one or both bromodomains (BD1 and/or BD2) of BET proteins, thereby displacing them from acetylated chromatin and suppressing oncogenic transcription. Despite their clinical progression, a comprehensive and comparative understanding of the precise atomic-level interactions between these inhibitors and their BET protein targets has remained incomplete. Elucidating these detailed molecular binding modes is crucial for rationalizing their mechanisms of action, interpreting clinical data, and informing the design of next-generation therapeutics with improved potency and selectivity.

To address this knowledge gap, we applied the structural biology approach to systematically investigate and compare the binding modes of these four clinical-stage inhibitors with a representative BET bromodomain. We focused on the second bromodomain (BD2) of human BRD2 (BRD2-BD2) as a model system. The recombinant BRD2-BD2 protein, fused to a glutathione S-transferase (GST) tag, was successfully expressed in *Escherichia coli* and purified to homogeneity using a multi-step chromatographic protocol. High-quality apo-protein crystals were obtained through extensive crystallization screening. The protein/inhibitor co-crystals were subsequently prepared using the soaking method, wherein pre-formed BRD2-BD2 crystals were incubated with high-concentration solutions of each individual inhibitor. X-ray diffraction datasets for all four complexes were collected at high resolutions, ranging from 1.4 to 1.5 Å. The three-dimensional structures were solved by molecular replacement and refined to excellent stereochemical statistics (**Table S1**).

The overall architecture of BRD2-BD2 in all complexes conforms to the canonical bromodomain fold, which is characterized by a left-handed bundle of four long α-helices (αZ, αA, αB, αC) interspersed with two short helices (α1 and αZ’) and connecting loop regions, most notably the ZA and BC loops (**Figure 1A, E, I, M**). In each refined model, the asymmetric unit contains a single BRD2-BD2 molecule bound to one inhibitor molecule in a 1:1 stoichiometry. All four small-molecule inhibitors are accommodated deep within the conserved hydrophobic Kac-binding pocket of BRD2-BD2, a cavity formed by the α1 helix, the ZA loop, the αZ’ helix, the αB helix, the BC loop, and the αC helix (**Figure 1B, F, J, N**).

**Figure 1.**
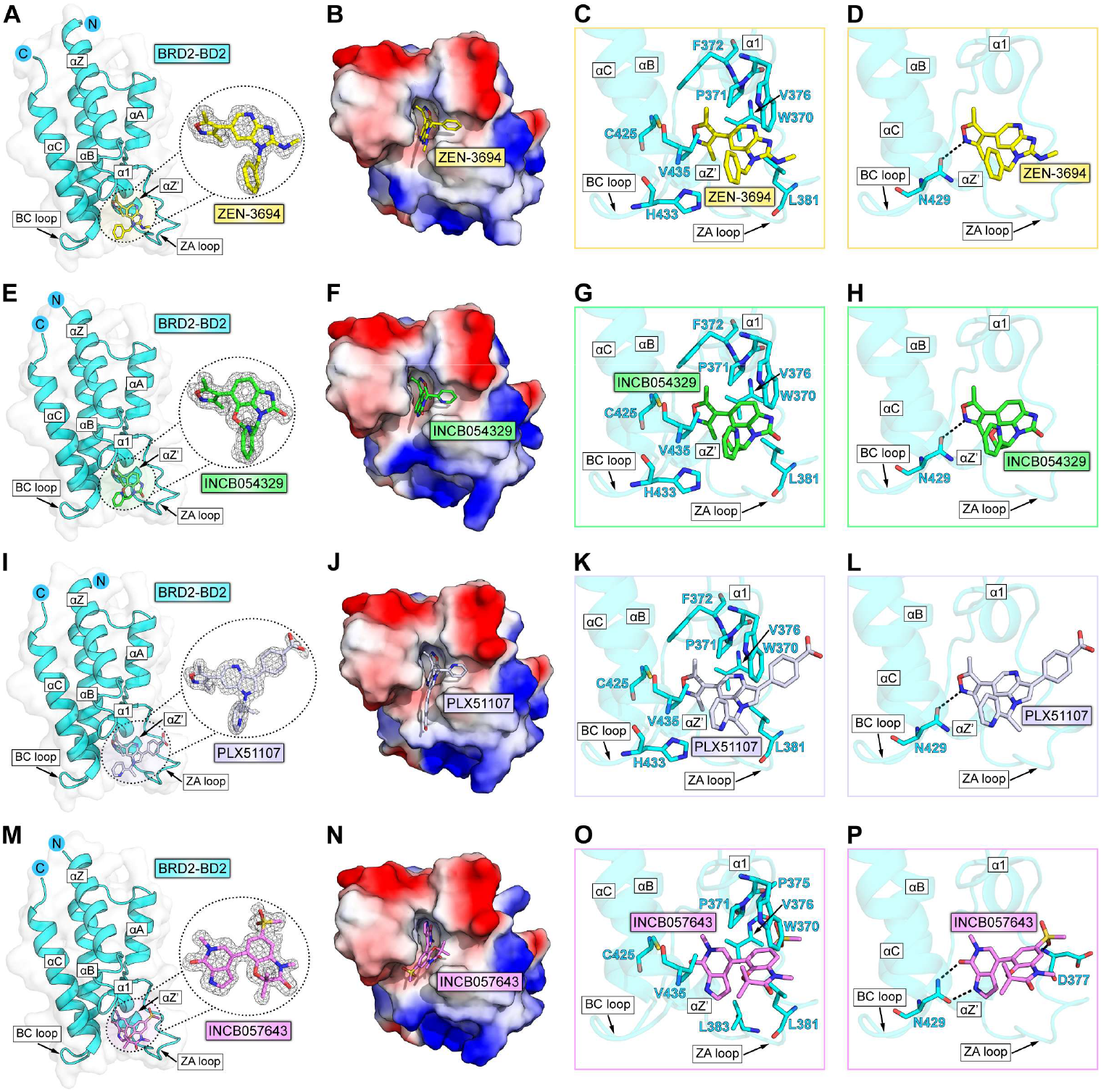
Structural analysis of human BRD2-BD2 in complex with four clinical-stage small-molecule inhibitors. (**A, E, I, M**) Overall views of BRD2-BD2 bound to ZEN-3694 (**A**), INCB054329 (**E**), PLX51107 (**I**), and INCB057643 (**M**). BRD2-BD2 is shown as a cyan cartoon. Inhibitors are depicted as sticks and colored yellow (ZEN-3694), green (INCB054329), blue-white (PLX51107), and violet (INCB057643). The corresponding 2|Fo|-|Fc| electron density maps (contoured at 1.0 σ) for the inhibitors are displayed within dashed circles. (**B, F, J, N**) Electrostatic surface representation of BRD2-BD2 in complex with ZEN-3694 (**B**), INCB054329 (**F**), PLX51107 (**J**), and INCB057643 (**N**). (**C, G, K, O**) Detailed views highlighting hydrophobic interactions between BRD2-BD2 and each inhibitor. (**D, H, L, P**) Detailed views highlighting hydrogen-bond interactions between BRD2-BD2 and each inhibitor. Hydrogen bonds are indicated by black dashed lines.

A detailed analysis of the interaction networks reveals consistent yet distinct binding patterns (**Figure 1C, D, G, H, K, L, O, P**). In the BRD2-BD2/ZEN-3694 complex, the inhibitor engages in extensive hydrophobic interactions with residues from multiple structural elements: W370, P371, and F372 on the α1 helix; V376 and L381 on the ZA loop; C425 on the αB helix; H433 on the BC loop; and V435 on the αC helix (**Figure 1C**). This hydrophobic network is coupled with a single hydrogen bond formed between ZEN-3694 and the side-chain of N429 located on the BC loop (**Figure 1D**).

Similarly, the structure of BRD2-BD2 in complex with INCB054329 shows the inhibitor forming a broad hydrophobic interface with an identical set of residues: W370, P371, F372, V376, L381, C425, H433, and V435 (**Figure 1G**). Analogous to ZEN-3694, INCB054329 also establishes one key hydrogen bond with the conserved N429 residue, underscoring the importance of this interaction (**Figure 1H**).

The binding mode of PLX51107 within the BRD2-BD2 pocket follows a closely related principle. It participates in intensive hydrophobic interactions with residues W370, P371, F372, V376, L381, C425, H433, and V435 (**Figure 1K**). Furthermore, PLX51107 forms a hydrogen bond to the side chain of N429, mirroring the polar interaction observed in the afore-mentioned complexes and contributing to binding stability (**Figure 1L**).

In contrast, the complex with INCB057643 exhibits a more elaborate interaction profile (**Figure 1O, P**). While it maintains a substantial hydrophobic network involving residues W370, P371, P375, V376, L381, L383, C425, and V435 (**Figure 1O**), its polar interaction signature is significantly enhanced. Notably, INCB057643 forms three hydrogen bonds with BRD2-BD2: one with the side chain of D377 on the ZA loop and two with the critical N429 residue on the BC loop (**Figure 1P**). This expanded hydrogen-bonding network indicates a potentially stronger and more specific association.

Collectively, these high-resolution structures demonstrate that the primary mechanism of binding for these diverse clinical inhibitors is through occupying the conserved hydrophobic Kac-recognition pocket of BRD2-BD2. A common and crucial auxiliary feature is the formation of at least one hydrogen bond with the side chain of N429 on the BC loop. Because this pocket is structurally and functionally dedicated to engaging acetylated lysine (12), these compounds act as direct competitive inhibitors, physically occluding the binding site and preventing the recruitment of BET proteins to acetylated substrates. This provides a clear molecular and structural rationale for their established transcriptional inhibitory effects.

From the structural comparison emerges a key observation with translational potential. Specifically, INCB057643, which was developed as a pharmacokinetically optimized successor (e.g., extended half-life) to INCB054329 (13), engages BRD2-BD2 via a more robust polar interaction network. We hypothesize that the additional hydrogen bonds, particularly those with D377 and N429, may contribute to a higher binding affinity and sustained target engagement, potentially underpinning its superior pharmacological profile *in vivo*. This finding illuminates a promising avenue for future structure-based drug design targeting the BET family proteins. It suggests that while maintaining optimal hydrophobic complementarity with the core of the Kac-binding pocket, the rational introduction or strengthening of polar interactions with key residues in the BC and ZA loop regions (e.g., D377, N429) could serve as a feasible strategy to enhance inhibitor potency and selectivity. This detailed structural framework not only deciphers the mechanism of existing clinical candidates but also provides a blueprint for the development of next-generation epigenetic therapies targeting the BET family.

## Data availability

Atomic coordinates and structure factors for the reported crystal structures have been deposited into the Protein Data Bank under accession codes XXXX, XXXX, XXXX and XXXX.

## Acknowledgements

We thank the Shanghai Synchrotron Radiation Facility of BL02U1 beamline (https://cstr.cn/31124.02.SSRF.BL02U1) for the assistance on X-ray diffraction data collection.

## Author contributions

G.L. and S.L. conceived the study and supervised the whole project. S.L. and Y.Y. conducted the majority of the experiments. S.L. and Y.Y. collected the datasets and solved the structures. X.C. assisted with the protein preparation. G.L., S.L. and Y.Y. wrote the manuscript.

