## Supporting Information for "Molecular basis of BET family protein inhibition by four clinical-stage small-molecule inhibitor drugs"

**This file includes:**

Materials and methods

Table S1

### Materials and methods

#### Plasmid construction

The DNA fragment encoding BRD2-BD2 was cloned into the pGEX-6p-1 expression vector. Briefly, primers were designed based on the genomic accession number (NM_001113182.3), incorporating BamH I and EcoR I restriction sites. PCR amplification was performed using PrimeSTAR Mix (Vazyme, China), and the resulting products were digested with BamH I and EcoR I. The purified fragments were then ligated into the prokaryotic expression vector pGEX-6p-1 via the corresponding restriction sites. Following standard transformation and plating, ampicillin-resistant colonies were picked. Plasmid extraction was carried out, and sequencing was performed to verify both the target gene and the junction regions, including the N-terminal GST-tag.

#### Gene expression and protein purification

After sequence verification, the recombinant plasmids were transformed into *E. coli* BL21 (DE3) cells. The BRD2-BD2 protein was expression under the induction of 500 μM isopropyl-β-D-thiogalactopyranoside (IPTG). For protein purification, the *E. coli* BL21 (DE3) cells were harvested and resuspended in the buffer containing 20 mM Tris-HCl (pH 8.0) and 150 mM NaCl. After lyse and centrifuge, the cleared lysate was incubated with pre-equilibrated GST resin (Cytiva). The bound GST-BRD2-BD2 fusion protein was eluted using the buffer containing 20 mM Tris-HCl (pH 8.0), 20 mM GSH, and 150 mM NaCl following extensive washing to remove the irrelative proteins. The GST tag was cleaved by PreScission Protease and removed. Finally, the target protein was collected and further purified by gel filtration using a Superdex™ 200 Increase 10/300 GL column (Cytiva).

**Crystallization**

For crystallization trials, BRD2-BD2 protein was concentrated and screened against commercial crystallization kits using the sitting-drop vapor diffusion method. Crystals were obtained by mixing 1 μL of protein solution (8 mg/mL) with an equal volume of reservoir solution at 8°C. The successful conditions were: 1. 0.09 M Halogens, 0.1 M Buffer System 2 (pH 7.5), 50% (v/v) Precipitant Mix 1; 2. 0.09 M NPS, 0.1 M Buffer System 2 (pH 7.5), 50% (v/v) Precipitant Mix 1; 3. 0.12 M Monosaccharides, 0.1 M Buffer System 2 (pH 7.5), 50% (v/v) Precipitant Mix 1.

**Soaking of small-molecule inhibitors into preformed crystals**

To obtain the BRD2-BD2/inhibitor complex structures, preformed BRD2-BD2 crystals were individually soaked in solutions containing the target inhibitors. Stock solutions of each compound (ZEN-3694, INCB054329, PLX51107, and INCB057643) were prepared at 50 mM in dimethyl sulfoxide (DMSO). Immediately prior to soaking, each stock was diluted with reservoir solution from the crystal growth condition to a final inhibitor concentration of 2.5 mM. Single crystals were then transferred into separate drops of the soaking solution and incubated at 8°C for approximately 12 hours to form the BRD2-BD2/inhibitor co-crystals.

**Data collection, processing and structure determination**

Prior to data collection, all of the crystals soaked with the four small-molecule inhibitors individually were cryo-protected through brief immersion in their respective reservoir solutions supplemented with 20% (v/v) glycerol, followed by rapid cooling in liquid nitrogen. The final X-ray diffraction data were collected at BL02U1 beamline at the Shanghai Synchrotron Radiation Facility in Shanghai, and subsequently processed by XDS (1) and scaled with Aimless (2) from CCP4 Software Suite (3).

The complex structures of BRD2-BD2/ZEN-3694, BRD2-BD2/INCB054329, BRD2-BD2/PLX51107, and BRD2-BD2/INCB057643 were determined through molecular replacement (4) in CCP4 suite (3) . Initial rigid-body refinement was carried out with Refmac5 (5) followed by iterative cycles of manual model rebuilding in Coot (6) and refinement in Phenix (7). Final statistics for data collection and structure refinement are summarized in **Table S1**. All structural figures were generated using PyMOL (https://pymol.org/).

**Table S1. Data collection and structure refinement statistics.**

| | BRD2-BD2/ZEN-3694 complex | BRD2-BD2/INCB054329 complex | BRD2-BD2/PLX51107 complex | BRD2-BD2/INCB057643 complex |
|---|---|---|---|---|
| **Data collection** | | | | |
| Space group | $P22_12_1$ | $P22_12_1$ | $P22_12_1$ | $P22_12_1$ |
| Cell dimensions | | | | |
| *a*, *b*, *c* (Å) | 31.99, 52.53, 72.03 | 31.98, 52.36, 72.17 | 32.18, 52.90, 72.58 | 32.02, 52.61, 72.02 |
| *α*, *β*, *γ* (°) | 90, 90, 90 | 90, 90, 90 | 90, 90, 90 | 90, 90, 90 |
| Wavelength (Å) | 0.97918 | 0.97918 | 0.97918 | 0.97918 |
| Resolution (Å) | 42.44-1.40 (1.42-1.40) | 42.38-1.40 (1.42-1.40) | 42.75-1.50 (1.53-1.50) | 42.48-1.50 (1.53-1.50) |
| Unique reflections | 24,432 (1,164) | 24,195 (1,189) | 20,258 (997) | 19,855 (979) |
| $R_{merge}$ | 0.040 (0.132) | 0.056 (0.361) | 0.063 (0.575) | 0.066 (0.542) |
| $I$/sig$I$ | 37.1 (12.2) | 19.6 (4.5) | 18.9 (4.0) | 21.8 (5.0) |
| Completeness (%) | 99.3 (98.4) | 98.3 (99.3) | 98.6 (100.0) | 98.4 (100.0) |
| Redundancy | 11.8 (9.2) | 7.2 (5.6) | 9.3 (8.5) | 11.5 (10.7) |
| **Refinement** | | | | |
| Resolution (Å) | 29.70-1.40 | 27.29-1.40 | 36.29-1.50 | 32.02-1.50 |
| No. reflections | 24,396 | 24,152 | 20,218 | 19,812 |
| $R_{work}/R_{free}$ | 0.162/0.168 | 0.168/0.168 | 0.171/0.182 | 0.173/0.178 |
| No. of atoms | | | | |
| Protein | 926 | 906 | 898 | 906 |
| Ligand/ion | 25 | 26 | 33 | 29 |
| Water | 172 | 142 | 132 | 128 |
| $B$-factors (Å$^2$) | | | | |

| | | | | |
|---|---|---|---|---|
| Protein | 14.7 | 15.2 | 19.2 | 16.8 |
| Ligand/ion | 18.7 | 15.2 | 27.5 | 18.0 |
| Water | 26.8 | 26.1 | 30.3 | 28.2 |
| R.m.s. deviations | | | | |
| Bond lengths (Å) | 0.009 | 0.009 | 0.008 | 0.008 |
| Bond angles (°) | 1.120 | 1.181 | 0.965 | 1.017 |
| Ramachandran plot (%) | | | | |
| Favored region | 99.10 | 99.07 | 99.07 | 100.00 |
| Allowed region | 0.90 | 0.93 | 0.93 | 0.00 |
| Outlier region | 0.00 | 0.00 | 0.00 | 0.00 |
| **PDB code** | XXXX | XXXX | XXXX | XXXX |

A single crystal was used to collect the data.

Values in parentheses are for the highest-resolution shell.
